# SARS-CoV-2 protein structure and sequence mutations: evolutionary analysis and effects on virus variants

**DOI:** 10.1101/2023.03.09.531961

**Authors:** Ugo Lomoio, Barbara Puccio, Giuseppe Tradigo, Pietro Hiram Guzzi, Pierangelo Veltri

## Abstract

Proteins sequence, structure, and function are related, so that any changes in the protein sequence may cause modifications in its structure and function. Thanks to the exponential growth of data availability, many studies have addressed different questions such as: (i) how structure evolves based on the sequence changes, (ii) how structure and function change over time. Computational experiments have contributed to the study of viral protein structures. For instance the Spike (S) protein has been investigated for its role in binding receptors and infection activity in COVID-19, hence the interest of scientific researchers in studying the effects of virus mutations due to sequence, structure and vaccination effects. Protein Contact Networks (PCNs) can be used for investigating protein structures to detect biological properties thorough network topology. We apply topological studies based on graph theory of the PCNs to compare the structural changes with sequence changes, and find that both node centrality and community extraction analysis play a relevant role in changes in protein stability and functionality caused by mutations. We compare the structural evolution to sequence changes and study mutations from a temporal perspective focusing on virus variants. We finally highlight a timeline correlation between Omicron variant identification and the vaccination campaign.

## Introduction

Comprehension of cellular processes requires the study of relations between the sequence of genes and the structure of encoded proteins [1, 2] through genomic and proteomic studies. From an evolutionary point of view, changes in gene sequences (e.g. single nucleotide mutations) may imply a modification of protein structure and thus phenotype changes. The evolutionary process limits some of the phenotype modifications on the basis of environmental constraints [1, 3].

The recent pandemic of the SARS-CoV-2 virus has led to the storage of a large amount of genomic and proteomic datasets enabling the molecular evolutionary analysis of functional data thereby boosting studies of the relations between the protein sequence structure and functions of the virus [4, 5]. The genome of SARS-CoV-2 contains 29.9 kilobase [6] and it has 14 functional open reading frames (ORFs), and multiple regions encoding: (i) four structural proteins, i.e., Spike (S), envelope (E), membrane (M) and nucleocapsid (N) protein; (ii) 16 nonstructural proteins (nsp1-nsp16) and (iii) accessory proteins [5, 7, 8].

Viruses undergo many mutations during the process of replication [9, 10]. Mutations can occur randomly due to errors in replication steps and to mutations on the structure of the viral proteins [11, 12]. These mutations may acquire an evolutionary advantage when they help the virus to evade the defence mechanism of the host’s immune system [13]. Many studies have identified mutations in SARS-CoV-2 [5, 14–16], and in the relation between virus mutations and capacity of escaping immune systems, e.g., relating the evolutionary action with the spread of the virus.

The accumulation of mutations may cause the insurgence of variants. A variant is a viral genome with one or more mutations causing changes in protein structure and virus characteristics. The World Health Organisation (WHO) and the European Center for Disease Control (ECDC) have the task of studying and assessing new evidence on variants. Sequenced samples of SARS-CoV-2 gathered from all countries, are analyzed by the WHO and ECDC with the aim of publishing information about variants and health related policies. Once a new variant is considered, the joint group ECDC/WHO updates the variants and the categories available at https://www.ecdc.europa.eu/en/covid-19/variants-concern and related data are stored in public databases [17–19]. Variants are organised into lineages, groups of closely related viruses with a common ancestor. Although mutations, i.e. a single change in the virus genome, occurs very frequently during virus replication, only a few modifications change the virus functionalities. A group of variants with similar genetic changes is designated by the WHO as a Variant Being Monitored (VBM), a Variant of Concern (VOC), or a Variant of Interest (VOI) due to shared attributes and characteristics that may require public health action [20, 21].

We focus on a subset of variants which cause modifications to the S protein structure, and with structural model description available at the exascale consortium database https://www.exscalate4cov.eu/ [22]. We study the impact of sequence changes on Spike protein structure by means of the Protein Contact Network (PCN) formalism [23–27]. PCNs are graphs whose nodes represent the *C* – *α* atoms of the backbone of proteins, while edges represent a relative spatial distance of between 4 and 8 angstroms among residues.Topological descriptors of PCNs, such as node centrality measures, are used to discover protein properties, even at the sub-molecular level [25, 26]. We investigate how sequence changes generate relevant changes in the structure. In order to do this, we calculate centrality measures for each node and their changes due to mutations. We also measure the structural differences between variants using the template modelling score (TM-SCORE) [28], which is a metric for assessing the topological similarity of protein structures. Moreover, for each considered PCN we extract communities by means of the Louvain algorithm [29] to see how many mutations belong to the same community. Then, we build two trees, one considering the difference in network descriptors and the second considering the distance in terms of TM-SCORE to clarify possible divergences between timeline evolutions of sequences and structure. Finally, we study the three different views for each variant: sequence, structure, and PCN network parameters. The obtained results have allowed us to put forward the following claims:

- The temporal evolution of the variants shows that both sequence and structure of Spike protein have undergone dramatic changes with the increase of vaccinations.
- Variants present changes in protein stability and functionality upon mutations in variants evidenced by changes in network structure.
- The analysis of centrality measures of nodes of PCN highlights that the Spike protein of the Omicron_1_ variant differs markedly from the Wild Type form and from Delta variant.
- Communities extracted with the Louvain algorithm seem to be correlated with the amino acid function in the protein. Some extracted communities once mapped in the three-dimensional structure of the protein, seem very similar to the well-known Receptor Binding Domain (RBD) and the N-Terminal Domain (NTD) of the Spike.
- Multiple mutations in Omicron variantsbelong to the same community and have the same to have the same outcome effect on protein function.
- Contact similarity rate between couples of PCNs variants, structural distance (TM-score), and sequence distance between variants are used to study the correlation between those three metrics of similarity of the protein.
- Amino acids polarity values *Pk_a_* for S protein in a variant shows a homogeneous variability pattern

Main contributions of the paper are: (i) the study of the correlation of evolution, sequence changes and structural changes in the Spike protein, in other words SARS-CoV-2 variations; (ii) the hypothesis that mutations of the Omicron variants which cause notable changes in the structure of the Omicron Spike may be related to the vaccination campaign.

## Materials and Methods

We here propose the analysis of sequence mutations and structural changes using the pipeline described in Fig. 1. The structures of the Spike protein of virus variants are used as input to the pipeline. As reported in Fig. 1, protein structures of Spike protein of selected variants are used as input for pipeline analysis. PDB files of the three-dimensional Spike structures are gathered from *escalate4cov* database [22]. Table 1 reports the variants used in the pipeline and the related mutations on Spike protein.

**Fig 1.**
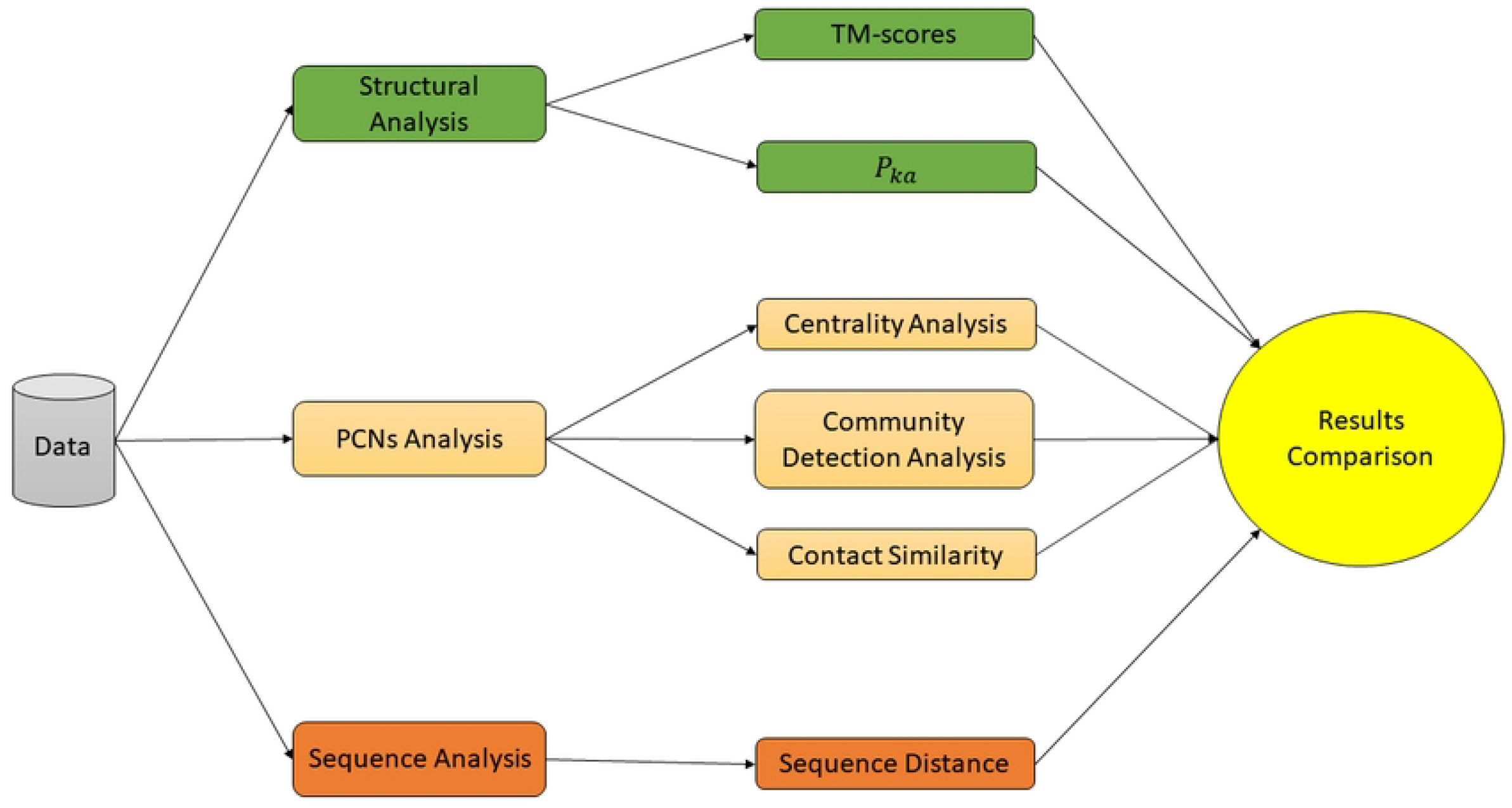
Analysis Workflow. We start from sequence and structure data of selected variants. We then analyse structure by calculating the structural differences by means of the TM-Score and the polarity (green boxes). Differences of the sequences are calculated to determine the distance among variants (orange box). Protein Contact Networks are used to analyse proteins on an intermediate scale. Finally, all obtained results are compared.

**Table 1.** SARS-CoV-2 Variants, lineage classification and mutations on the Spike Protein sequence.

| Variant | Lineage | Mutations on Spike Protein |
| --- | --- | --- |
| Alpha $\alpha$ | B.1.1.7 | HV69-70Δ, Y144Δ, N501Y, A570D, D614G, P681H, T716I, S982, D1118H |
| Beta $\beta$ | B.1.351 | D80A, D215G, 241-243Δ, R246I, K417N, E484K, N501Y, D614G, A701V |
| Omicron<sub>1</sub> $o_1$ | BA.1 | A67V, HV69-70Δ, T95I, G142D, V143Δ, YY144-145Δ, N211Δ, L212I, ins214EPE, G339D, S371L, S373P, S375F, K417N, N440K, G446S, S477N, T478K, E484A, Q493R, G496S, Q498R, N501Y, Y505H, T547K, D614G, H655Y, N679K, P681H, N764K, D796Y, N856K, Q954H, N969K, L981F |
| Omicron<sub>5</sub> $o_5$ | BA.5 | Δ25-27, HV69-70Δ, G142D, V213G, G339D, S371F, S373P, S375F, T376A, D405N, R408S, K417N, N440K, L452R, S477N, T478K, E484A, F486V, Q498R, N501Y, Y505H, D614G, H655Y, N679K, P681H, N764K, D796Y, Q954H, N969K |
| Gamma $\gamma$ | P.1 | D138Y, R190S, K417T, E484K, N501Y, D614G, H655Y, T1027I |
| Zeta $\zeta$ | P.2 | E484K, D614G, V1176F |
| Delta $\delta$ | B.1.617.2 | EE156-157Δ, R158G, L452R, T478K, D614G, P681R, D950N |
| Kappa $\kappa$ | B.1.617.1 | E154K, E484Q, D614G, P681R, Q1071H |
| Epsilon $\epsilon$ | B.1.427 | S13I, W152C, L452R, D614G |
| Eta $\eta$ | B.1.525 | Q52R, A67V, HV69-70Δ, Y144Δ, E484K, D614G, Q677H, F888L |
| Iota<sub>1</sub> $\iota_1$ | B.1.526 | L5F, T95I, D253G, S477N, D614G |
| Iota<sub>2</sub> $\iota_2$ | B.1.526 | L5F, T95I, D253G, E484K, D614G |
| Ihu | B.1.640.1 | E96Q, CNDPFLGVY136-144Δ, R190S, I210T, R346S, N394S, Y449N, F490R, N501Y, D614G, P681H, T859N, D936H, K1191N |

For each Spike protein we compute a corresponding PCN using the PCN-Miner tool [30]. Each node of the PCN corresponds to a single amino acidof the proteins, while an edge connects two nodes whose spatial distance is comprised between 4 and 8 angstroms. For each node of the PCN a set of centrality measures is evaluated. The measures and their definition are reported in the following:

- *Betweenness Centrality measure*: given a node *i,* this value measures how much the node (i.e., amino acid) influences communication and serves as a bridge from one part of the graph (i.e., part of the represented protein) and another.

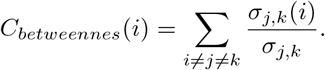

where, *σ_j,k_* indicates the number of shortest paths from node *j* to node *k* and *σ_j,k_*(*i*) is the shortest path which includes *i*.
- *Degree Centrality measure*: given a node i, this measures the normalised degree of the node i, i.e. the number of connections of the node. Nodes (amino acids) with a high degree centrality are considered hubs in the network (i.e., the protein) and have a crucial role in the network communication. The degree centrality of a node *i* is computed as follows;

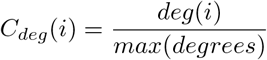
- *Eigenvector centrality measure*: given a generic node i, this measures the transitive influence of the node. Such a measure allows researchers to discover a possible interaction between the node *i* and a generic node *j,* and it also takes the eigenvector centrality score of the considered node into account. Nodes that interact with node *i* and have a high eigenvector centrality score contribute to the eigenvector centrality value of the *i* node. If node *i* has a high eigenvector centrality value this means that the node is connected to many nodes which also have high scores. The eigenvector centrality of a node *i* is equal to the *i-th* element of vector *x* defined as follows:

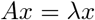
- *Closeness centrality measure*: given a node *i*, this measure how close the node (amino acid) is to all the other nodes in the graph (protein) and it is evaluated as follows:

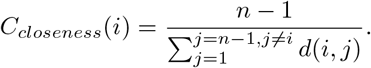

where *d*(*i*,*j*) is the shortest distance between *i* and *j*.
- *Katz centrality measure*: given a generic node i, this measures the relative degree of influence of the node (amino acid) in the graph (protein) and is obtained as follows:

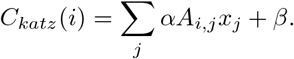

where *α* and *β* are parameters that indicate respectively: (i) the attenuation factor and (ii) the weight attributed to the immediate neighborhoods of each node.

We measure the centrality values of all the PCN nodes and compare both the overall changes (i.e. averaging the centrality values) and local changes (i.e. changes in the centrality values of mutated residues). We depict these values by using boxplots and radar plots. Boxplots are associated to all variants.Radar plots are used to represent centrality values of mutated nodes in the Spike variants. Finally, the obtained centrality measures are mapped onto the real protein structure using PCN-Miner. A t-test [31] on the variants centrality distribution is used to evaluate the significance of any changes.

Community detection analysis has been performed with the Louvain algorithm to study the relation between virus mutations and nodes communities in PCNs. For each variant, we plot its mutations inside the communities to graphically show in which community most of the mutations end up. This allows us to identify a pattern in mutation distributions of the Louvain communities. In the case of a high percentage of mutations belonging to the same community, we can claim that the corresponding mutations share similar effects on protein functionality.

We computed TM-scores [32] between pairs of Spike proteins of two different variants. The TM-score quantify the structural similarity between proteins. The US-align (Universal Structural alignment) software [33] has been used. Thus, sequence distance and structural distance between the Spike variants are measured and plotted on Fig. 11. Contact similarity between PCNs variants, defined as the percentage of contacts/non-contacts shared between two PCNs, is also evaluated and plotted.

**Fig 2.**
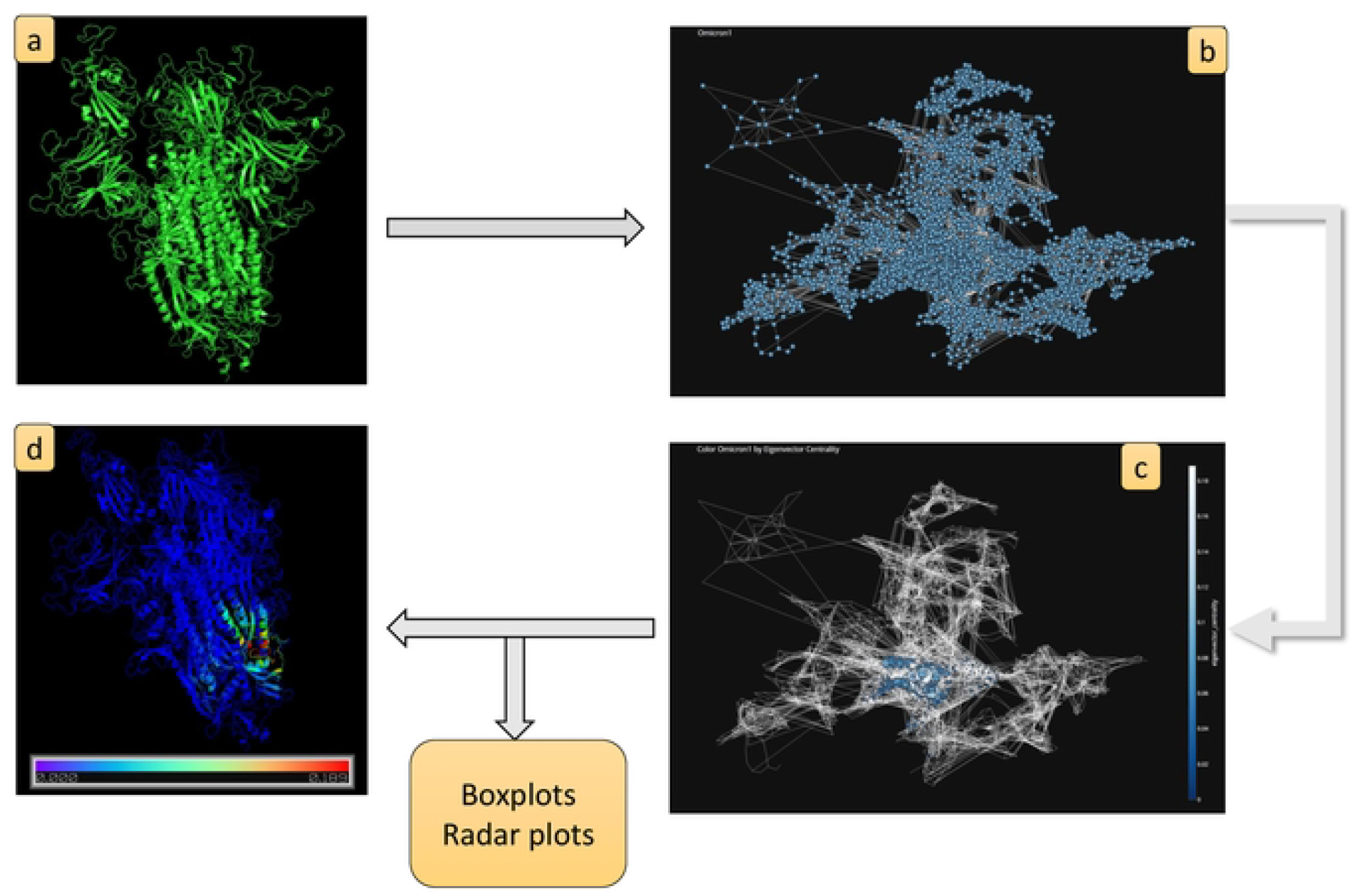
Centrality analysis workflow: a) Start with a PDB file (Spike Omicron_1_ variant in this example); b) compute the corresponding PCN; c) apply a node centrality measure algorithm (in this example eigenvector centrality); d) map nodes centrality values directly on the protein structure.

**Fig 3.**
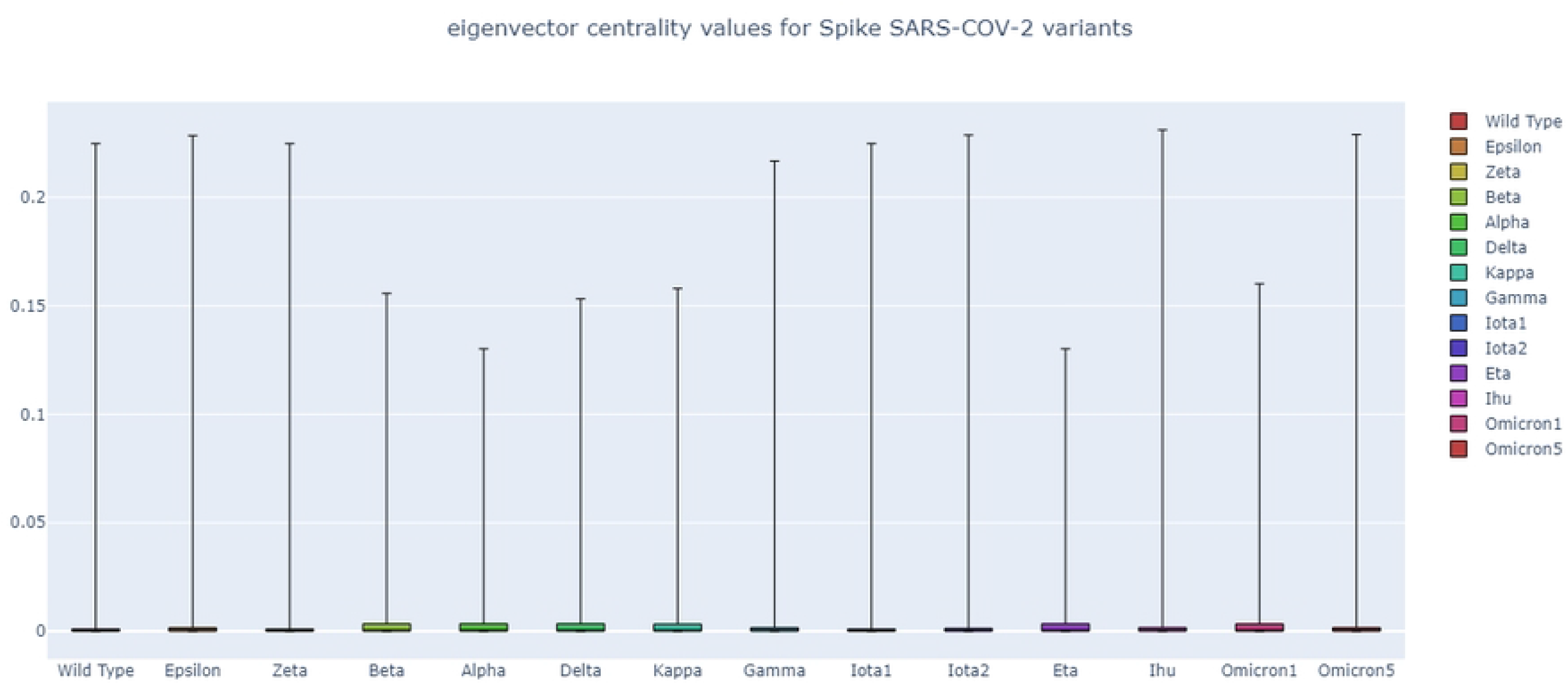
Node eigenvector centrality values for all variants. We report changes in node eigenvector centrality values in all variants. Each box represents nodes eigenvector centrality values of a specific variant. Variants are ordered on the basis of the date of their first appearance.

**Fig 4.**
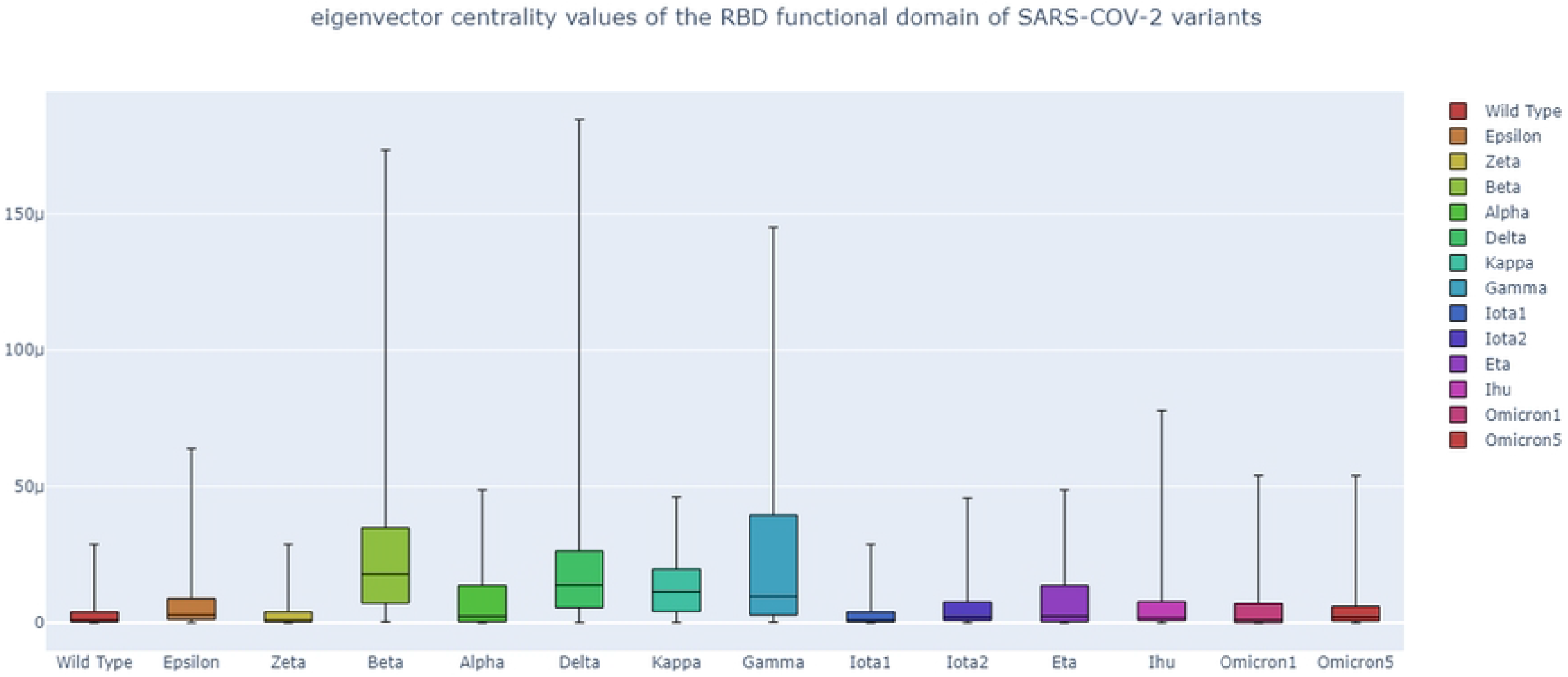
Node eigenvector centrality values in the RBD functional domain of all variants. We report changes in node eigenvector centrality values in the RBD functional domains of all variants. Each box represents RBD nodes eigenvector centrality values of a variant. Variants are ordered from the date of their first appearance.

**Fig 5.**
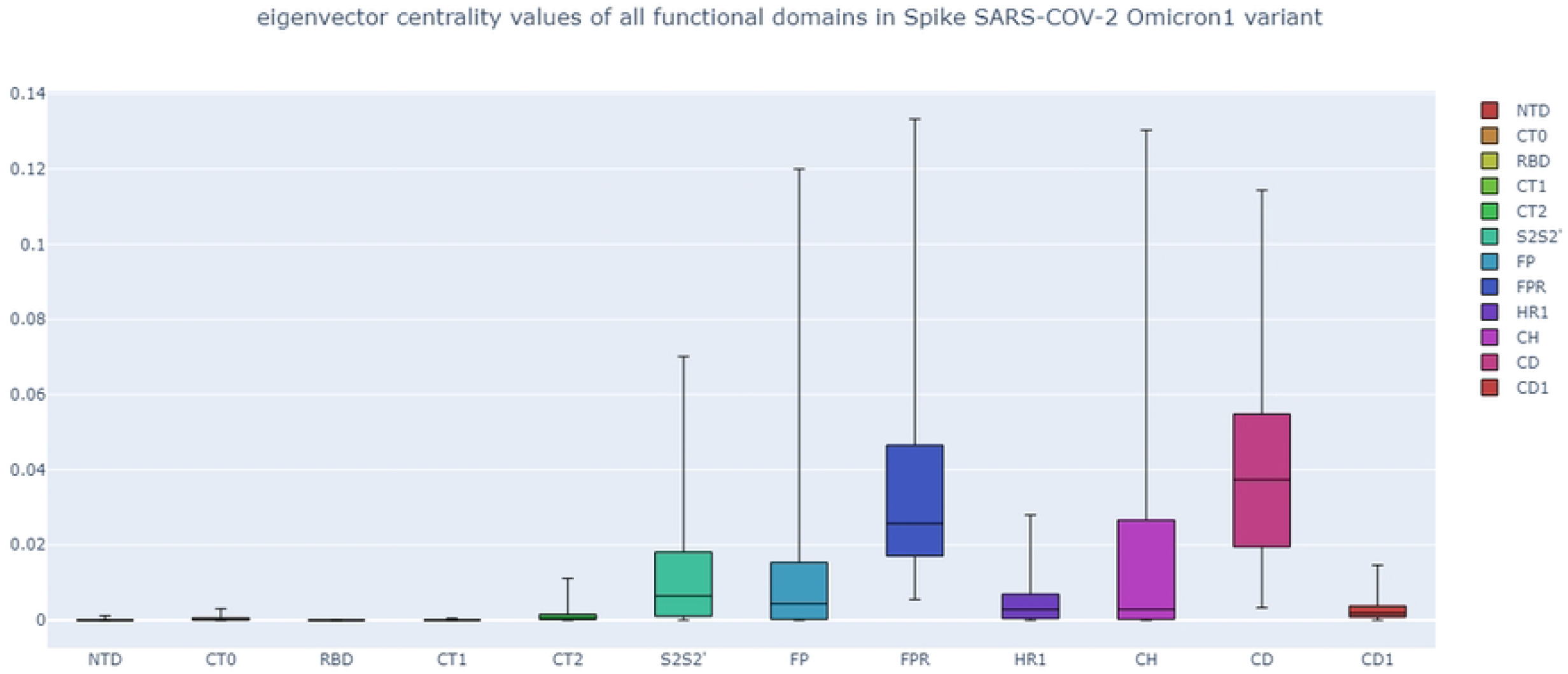
Node eigenvector centrality distribution in all functional domains of the Omicron_1_ variant. We report the distribution of the node eigenvector centrality values distribution in all functional domains of a single variant. Each box represents functional domain nodes eigenvector centrality values. As can be seen, in the Omicron_1_ variant the fusion peptide region (FPR) and the connecting domain (CD) have higher eigenvector centrality values. The Omicron_1_ variant presents a mutation in the first amino acid of this FPR domain, the N856K of the Omicron_1_ variant

**Fig 6.**
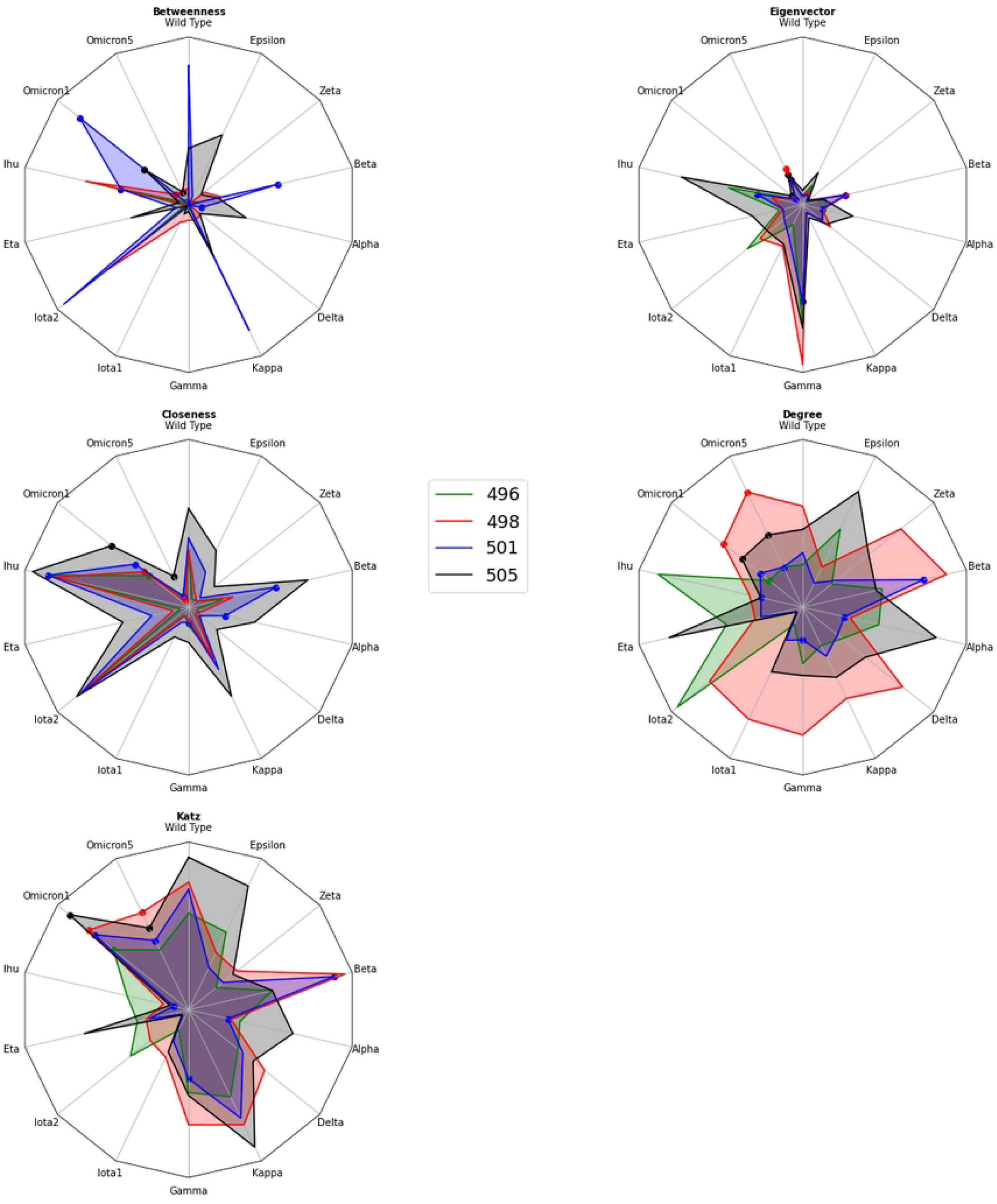
Radar plots of centrality values in RBDs mutated nodes from amino acid number 496 to 505. Colours are used to identify nodes 496 (red), 498 (green), 501 (blue), and 505 (black). Mutation in situ is indicated by a single point in the figure. From the upper corner representing Wild Type, variants are ordered clockwise from the date of their first appearance. The Figure shows that mutations also have an impact on the centrality of non-mutated points in the RBD domain.

**Fig 7.**
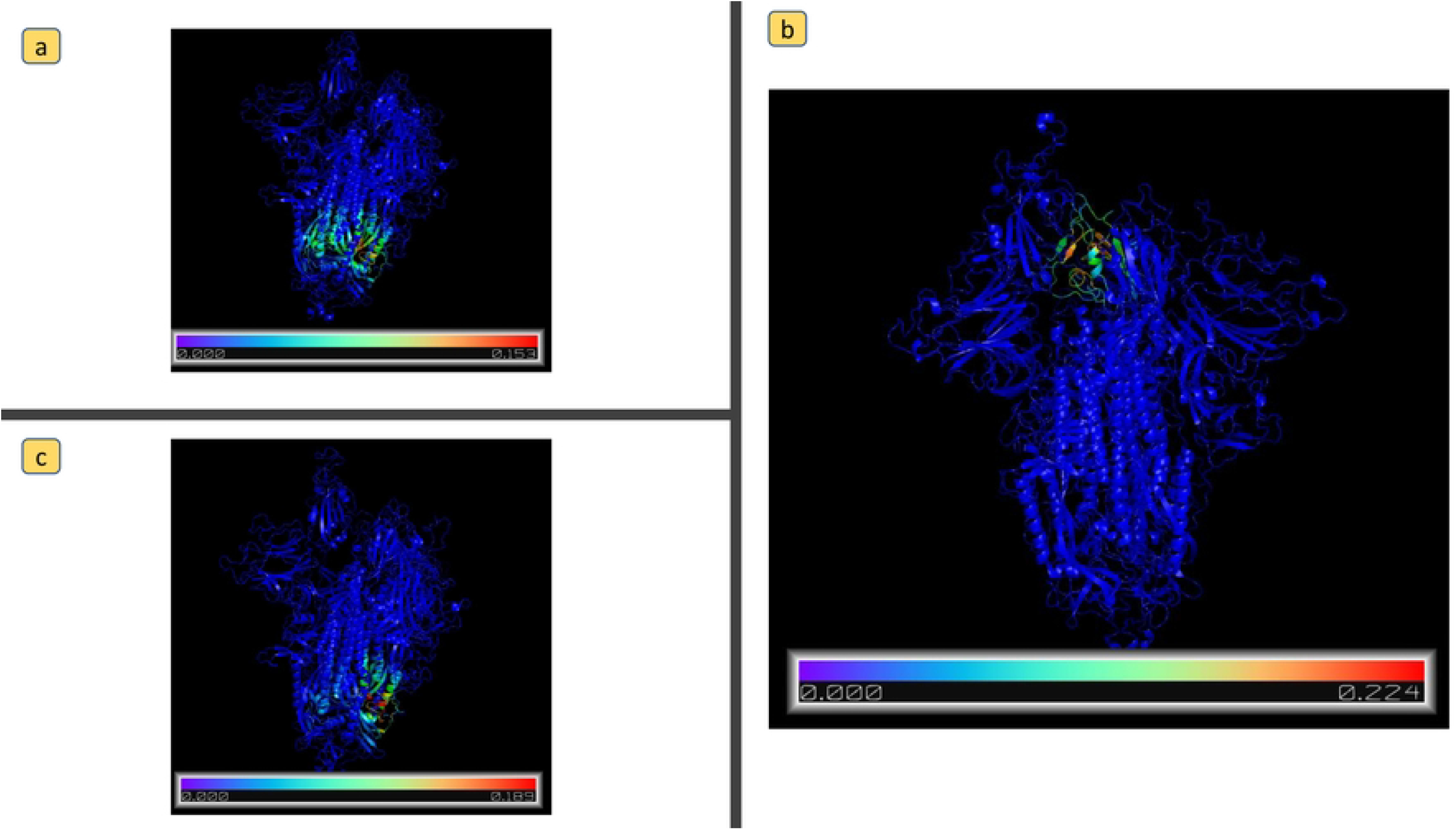
Amino acids eigenvector centrality values mapped on the protein structure of the following Spike variants: a) Omicron_1_; b) Wild Type; c) Delta.

**Fig 8.**
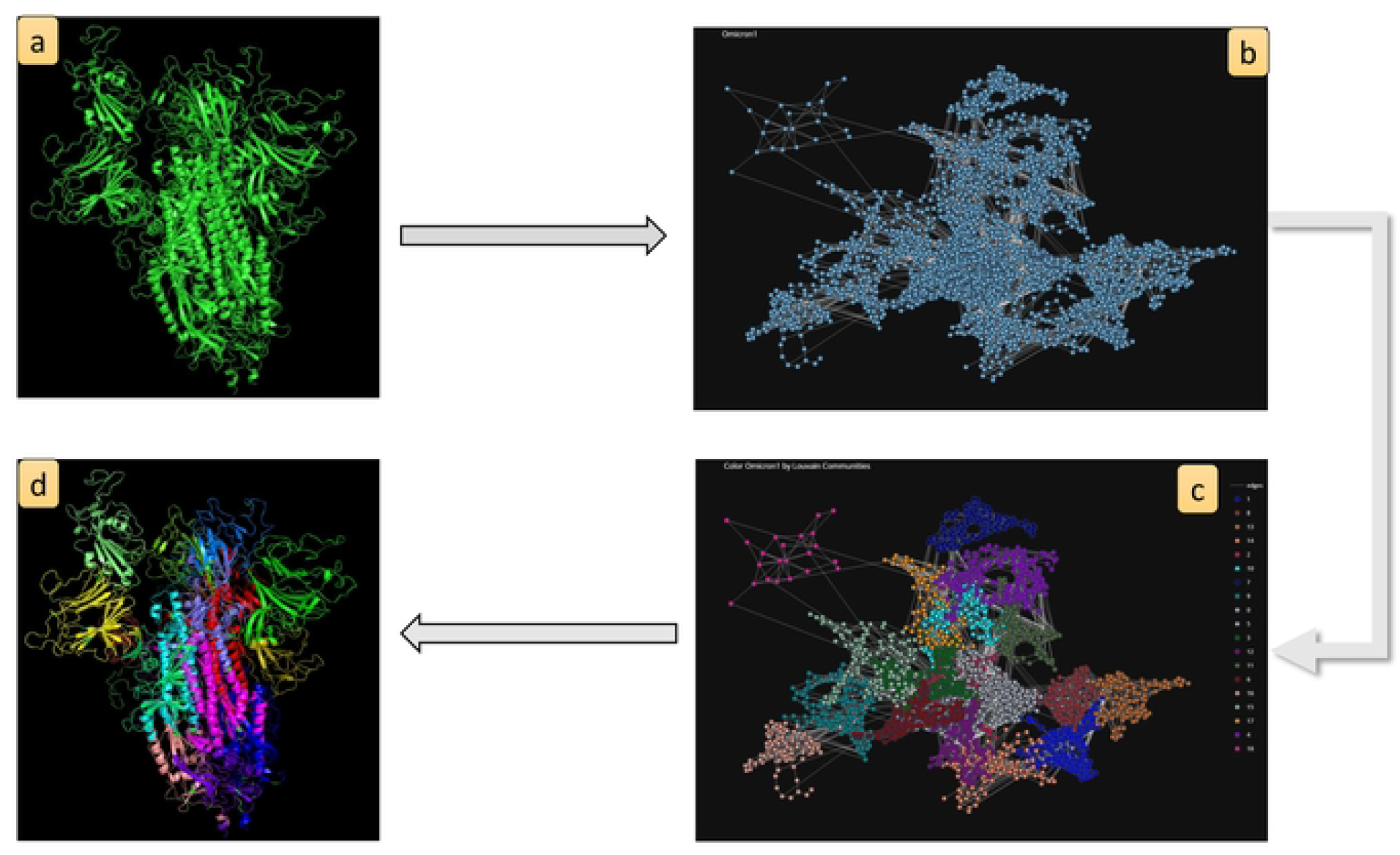
Community detection analysis workflow. Starting from a PDB file, in this example the SARS-CoV2 Omicron_1_ Spike variant, we compute the corresponding PCN and apply the Louvain community detection algorithm. In this way, we can map communities directly on the protein structure or visualize mutations inside the communities.

**Fig 9.**
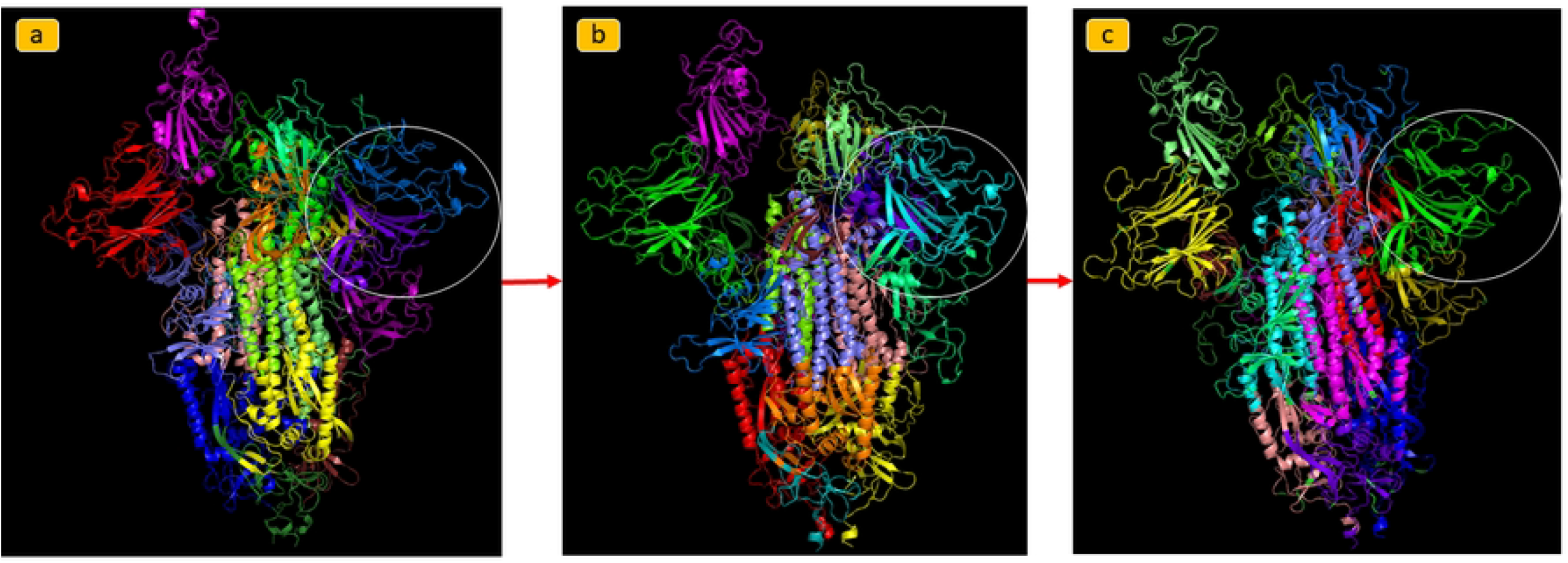
Community detection analysis comparing Spike of the Wild Type, Delta, and Omicron_1_ variant. Communities mapped directly on the protein structure of the a) Wild Type, b) Delta, and c) Omicron_1_ variant to visualize functional domains predicted by the Louvain algorithm. The Louvain founds 19 communities in the Spike Wild Type, Delta, and Omicron_1_ variants.

**Fig 10.**
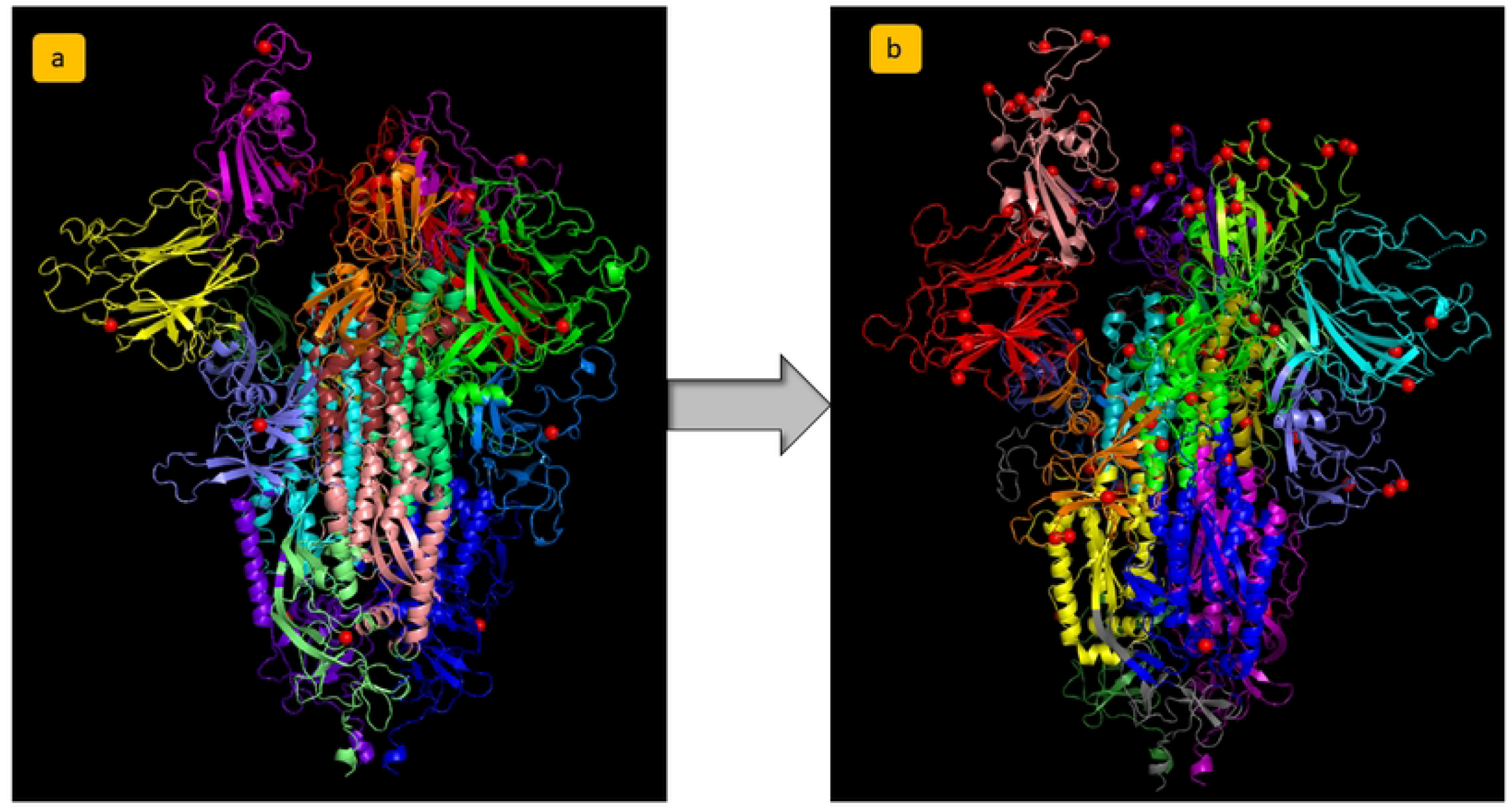
Community detection analysis comparison between a) Delta and b) Omicron_1_. Visualization of the mutations and the communities on the protein structure. Mutated residues are displayed as red spheres. In Omicron_1_, and in Omicron_5_, the mutations seem to fall inside certain communities with the same function in the Spike Protein. As can be seen in b) part of the figure, Omicron_1_ has more than ten mutations that fall inside the same community which, moreover, seems to correspond to the RDB functional domain.

**Fig 11.**
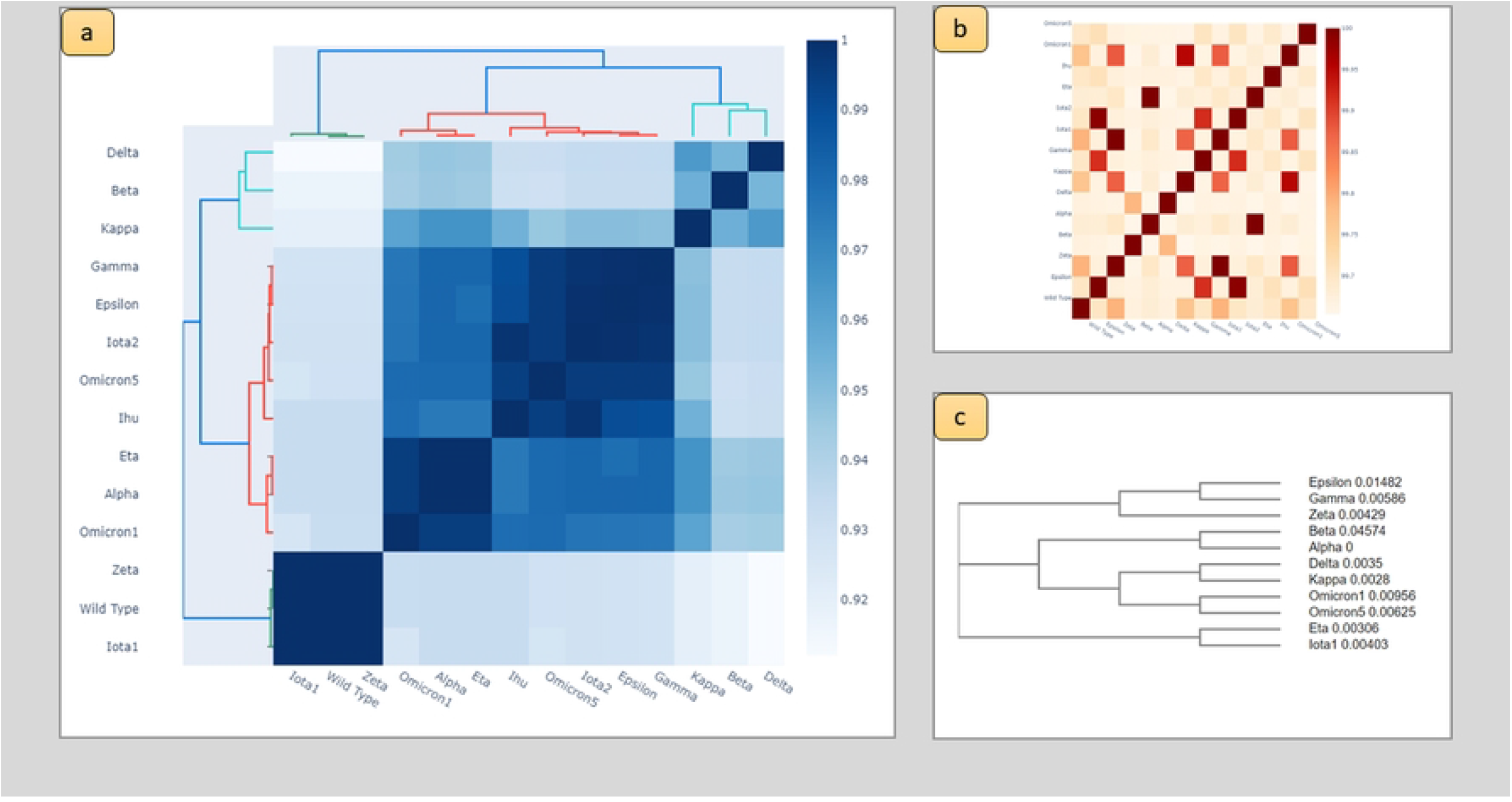
Structural, Sequence and Network similarity analysis results: a) Heatmap representing the structural similarity of variants computed by TM-scores; b) Heatmap representing the similarity between PCN from variants; c) sequence similarity represented by a Phylogenetic tree obtained from Clustal Omega. The figure evidences that there is no a direct correlation between sequence changes and structural modification. For instance, Iota1, and Zeta variants have dissimilar sequences but very similar structures.

Finally, for each variant we computed the acid dissociation constant *Pk_a_* for each amino acid of the analysed proteins using the PROPKA3 web server [34]. Given a node, *Pk_a_* of the node stands for the −*log*_10_*K_a_*, where *K_a_* is the acid dissociation constant (*K_a_*) that measures amino acid acidity or basicity.

### Implementation

The methods described above have been developed in the JupyterLab development environment [35] with Python language version 3 and also by: (i) plotly and matplotlib libraries to draw figures and (ii) propka [34] library to estimate the whole *Pk_a_* of the proteins. The PCN-Miner [30] tool is used to compute PCNs, node centralities, and Louvain algorithms to extract communities [29] for the Spike variants. Finally, PyMOL library is used to read, visualize and modify PDB files. The CUPSAT software [36] is used to predict changes in protein stability caused by a particular mutation of the Wild Type form. The prediction model uses amino acid atom potentials and torsion angle distribution to assess the amino acid environment of the mutation site. We use US Align [33] to perform a sequence-independent alignment based on the structural similarity of the variants and compute the TM-score.

## Results and Discussion

Starting from the PDB files of the SARS-CoV-2 Spike protein variants we computed the PCNs, and evaluated centrality measures as reported in Fig. 2. We used PCN-Miner to define PCNs and performed measurements intra PCN and betweein PCNs. Centrality measures are represented as boxplots in Figs. 3, 4 and 5 focusing on: (i) the comparison of the whole protein in all variants, and (ii) the RBD domain of Omicron variant. Fig. 3 reports boxplots regarding eigenvector centrality. Fig. 4 reports eigenvector centrality values for the RBD domain for all the variants. In Fig. 5, we set a variant and analysed each domain regarding node eigenvector centrality. We further compared the centrality values of the mutation sites to highlight possible differences on the same variant sites. Variants have been ordered in a clockwise fashion on the basis of their detection on a radar plot from the Wild Type (see Fig. 6). Measurement results and boxplots are available at https://github.com/UgoLomoio/SARSCoV2_variants_PCN.

Changes in eigenvector centrality evidence protein instability [23]. According to [37], mutations in Omicron_1_ and Delta variants cause Spike protein instability. We note that for the Wild Type virus, amino acids of the RBD domain present the highest value for eigenvector centrality measure. The Omicron_1_ variant has 15 mutations in the RBD domain, 14 of which can be related to protein instability, whereas N501Y mutation site is the only Omicron_1_ mutation in the RBD that increases protein stability). In the Omicron_1_ PCN, the decreased protein stability caused by mutations in RBDs causes a decrease in the RBD eigenvector centrality values. These results are reported in Fig. 7. We measured the significance of the changes in centrality measures by comparing average values by means of a t-test. We compared the average centrality of all the nodes and the average centrality of nodes corresponding to mutations (see Tables 2 and 3). We found a significant variation of Katz centrality measures between Omicron_1_ variant and all the other Spike variants (Wild Type included). Significant changes in betweenness centrality and degree centrality values have also been reported, where the significance is valued by using t-test analyses. Comparisons of measures for all the considered variants can be found at https://github.com/UgoLomoio/SARSCoV2_variants_PCN.

**Table 2.** T-test p-values for the comparison of average node centrality values of mutated nodes in the Omicron_1_ variant only.

| Variants couples | p-value EC | p-value BC | p-value KC | p-value DC | p-value CC |
| --- | --- | --- | --- | --- | --- |
| Omicron<sub>1</sub> vs Wild Type | 0.655 | 0.353 | 1.148e-10 | 0.181 | 0.091 |
| Omicron<sub>1</sub> vs Epsilon | 0.627 | 0.127 | 8.110e-04 | 0.820 | 0.648 |
| Omicron<sub>1</sub> vs Zeta | 0.655 | 0.353 | 1.148e-10 | 0.181 | 0.091 |
| Omicron<sub>1</sub> vs Beta | 0.1 | 0.00000077 | 1.442e-03 | 0.001 | 0.022 |
| Omicron<sub>1</sub> vs Alpha | 0.551 | 0.157 | 1.705e-01 | 0.410 | 0.763 |
| Omicron<sub>1</sub> vs Delta | 0.22 | 0.00002 | 2.619e-02 | 0.004 | 0.139 |
| Omicron<sub>1</sub> vs Kappa | 0.818 | 0.004 | 1.417e-01 | 0.380 | 0.934 |
| Omicron<sub>1</sub> vs Gamma | 0.674 | 0.011 | 1.053e-04 | 0.888 | 0.923 |
| Omicron<sub>1</sub> vs Iota<sub>1</sub> | 0.671 | 0.314 | 5.270e-11 | 0.242 | 0.099 |
| Omicron<sub>1</sub> vs Iota<sub>2</sub> | 0.695 | 0.310 | 6.346e-03 | 0.663 | 0.825 |
| Omicron<sub>1</sub> vs Eta | 0.862 | 0.166 | 3.758e-01 | 0.880 | 0.870 |
| Omicron<sub>1</sub> vs Ihu | 0.008 | 0.105 | 8.761e-10 | 0.014 | 0.088 |
| Omicron<sub>1</sub> vs Omicron<sub>5</sub> | 0.397 | 0.226 | 6.490e-04 | 0.027 | 0.690 |

**Table 3.** T-test p-values for the comparison of average node centrality values of all nodes in the Omicron_1_ variant. Values less than 0.01 after correction for multiple test were considered significant.

| Variants couples | p-value EC | p-value BC | p-value KC | p-value DC | p-value CC |
| --- | --- | --- | --- | --- | --- |
| Omicron<sub>1</sub> vs Wild Type | 0.725 | 3.293e-31 | 1.340e-119 | 1.991e-18 | 0.00002 |
| Omicron<sub>1</sub> vs Epsilon | 0.925 | 1.097e-02 | 2.378e-45 | 1.323e-03 | 0.0597 |
| Omicron<sub>1</sub> vs Zeta | 0.725 | 3.293e-31 | 1.340e-119 | 1.991e-18 | 0.00002 |
| Omicron<sub>1</sub> vs Beta | 0.192 | 4.850e-13 | 1.120e-23 | 2.037e-07 | 0.067 |
| Omicron<sub>1</sub> vs Alpha | 0.591 | 1.865e-04 | 1.328e-01 | 1.334e-11 | 0.555 |
| Omicron<sub>1</sub> vs Delta | 0.027 | 4.552e-12 | 5.673e-23 | 3.599e-08 | 0.050 |
| Omicron<sub>1</sub> vs Kappa | 0.609 | 1.067e-05 | 7.881e-19 | 1.423e-07 | 0.266 |
| Omicron<sub>1</sub> vs Gamma | 0.916 | 7.393e-01 | 3.139e-57 | 2.852e-04 | 0.773 |
| Omicron<sub>1</sub> vs Iota<sub>1</sub> | 0.725 | 3.293e-31 | 1.340e-119 | 1.991e-18 | 0.00002 |
| Omicron<sub>1</sub> vs Iota<sub>2</sub> | 0.930 | 7.954e-07 | 2.450e-23 | 6.653e-04 | 0.405 |
| Omicron<sub>1</sub> vs Eta | 0.591 | 1.865e-04 | 1.328e-01 | 1.334e-11 | 0.555 |
| Omicron<sub>1</sub> vs Ihu | 0.643 | 2.318e-04 | 2.132e-34 | 4.292e-04 | 0.519 |
| Omicron<sub>1</sub> vs Omicron<sub>5</sub> | 0.527 | 3.886e-05 | 3.349e-20 | 1.644e-05 | 0.946 |

We also studied how mutation sites can be topologically related. Studying PCNs, we focus on how mutation sites occur in the same communities identified by subgraphs characterised by higher modularity representing potentially related nodes. We performed a community detection analysis for each PCN (each Spike variant) using the Louvain algorithm [29]. We noted that, in most cases, the obtained Louvain communities correspond to the functional domains of the Spike protein. Fig. 8 explains the community detection analysis workflow with the Spike Omicron_1_ variant as an example. Communities are then mapped onto the three-dimensional protein structure to show how they relate to different substructures. Mutations that fall inside the same community, have the same effect on the protein function [24]. The resulting communities, extracted from the studied dataset, can be found at https://github.com/UgoLomoio/SARSCoV2_variants_PCN/tree/main/Figures/Mutations_In_Communities_html. Moreover Figs. 9 and 10 provide community detection analysis results. The obtained communities are similar to the functional domain of the protein. Moreover, Fig. 9 reports a comparison of mutated nodes distribution inside Louvain communities between Delta, Omicron_1_, and Omicron_5_ variants, where mutated nodes are in red. As shown in Fig. 10, Omicron_1_, and Omicron_5_ RBD mutations fall inside a single community, meaning that these mutations may have similar effect on changes in Spike protein function. Note that such a distribution of multiple mutations in the same Louvain community was not found in any previous identified virus variant (such as the Delta). Table 4 summarises results of Louvain community analysis.

**Table 4.** Summary of communities and their mutations. The UM (UnModelled) is a special community whose group amino acids belong to regions without a determined structure.

| Variant | n.° communities | n.° mutations | mutations for each community |
| --- | --- | --- | --- |
| Wild Type | 22 | 0 |  |
| Epsilon | 22 | 4 | UM: 1, Community 0: 1, Community 4: 1, Community 16: 1 |
| Zeta | 19 | 3 | Community 9: 1, Community 8: 1, UM: 1 |
| Beta | 21 | 8 | Community 4: 3, Community 15: 3, Community 11: 1, Community 7: 1 |
| Alpha | 22 | 7 | Community 10: 1, Community 11: 1, Community 12: 2, Community 18: 2, Community 6: 1 |
| Delta | 22 | 6 | Community 1: 3, Community 15: 2, Community 0: 1 |
| Kappa | 23 | 5 | Community 1: 2, Community 15: 2, Community 5: 1 |
| Gamma | 19 | 8 | Community 3: 2, Community 6: 2, Community 16: 1, Community 10: 2, Community 4: 1 |
| Iota1 | 21 | 5 | UM: 1, Community 2: 2, Community 15: 1, Community 9: 1 |
| Iota2 | 21 | 5 | UM: 1, Community 0: 2, Community 15: 1, Community 8: 1 |
| Eta | 24 | 6 | Community 19: 2, Community 1: 1, Community 12: 2, Community 10: 1 |
| Ihu | 24 | 13 | Community 15: 1, Community 1: 4, Community 7: 3, Community 12: 2, Community 0: 2, UM: 1 |
| Omicron<sub>1</sub> | 20 | 30 | Community 2: 4, Community 5: 15, Community 7: 1, Community 9: 4, Community 13: 4, Community 0: 2 |
| Omicron<sub>5</sub> | 19 | 27 | Community 2: 10, Community 7: 8, Community 17: 1, Community 3: 4, Community 13: 2, Community 0: 2 |

Protein structures were analyzed by means of TM-scores, structural distance, and PCN Contact similarity between Spike variants. Sequence distances between variants were used to construct a phylogenetic tree. Fig. 11 reports the comparison of such measures.

For each variant, we computed, the acid dissociation constant (*PK_a_)* of each amino acid to measure its acidity or basicity inside the protein. Fig 12 reports *PK_a_* values for Omicron_1_, Delta, and Wild Type. Note that the Delta variant has a single mutation in the RBD, i.e., the L452R. This mutation causes a negative *PK_a_* region that is not present in the Wild Type or Omicron_1_ variant (that has 15 RBDs mutations). Additionally, from amino acid 452 to 768, the Delta variant has more negative *PK_a_* regions which may be of interest for future studies.

**Fig 12.**
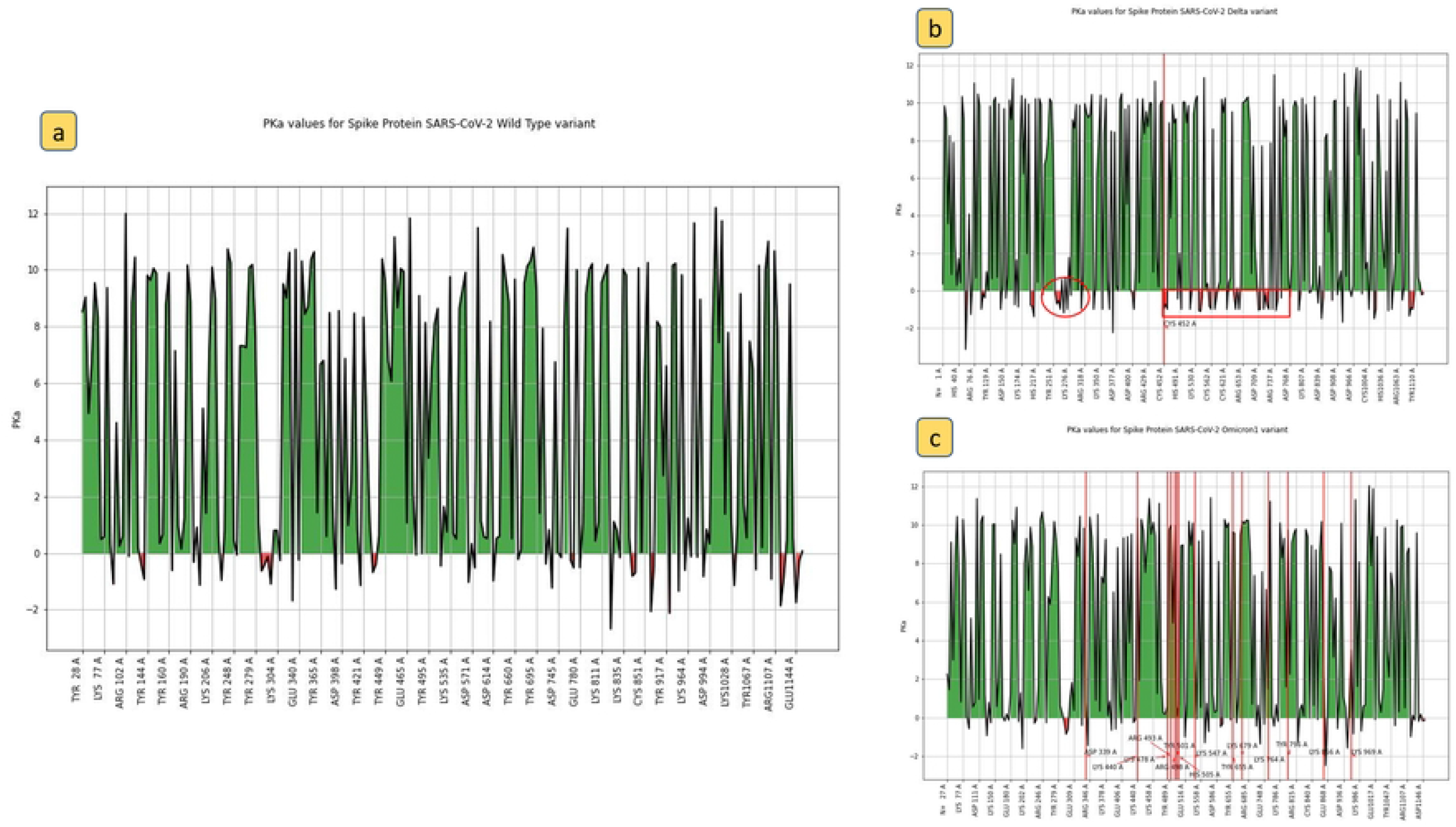
*PK_a_* values for a) Wild Type; b) Delta variant; c) Omicron_1_ variant.

In conclusion, by integrating multiple data sources [15], we analysed the mutation pattern of SARS-CoV-2 genome, focusing on the mutations of the S protein. We studied a set of variants, including the VOC and other ones. We analysed the impact of these mutations on the structure of the S protein by the PCN formalism [27].

We found that mutant variants have a lower average value of closeness centrality, thus these structures are less able to propagate signals. Our results show that Omicron variants clearly differ from the others. All the considered parameters confirm that there is a remarkable change in centrality values. In particular, the difference in terms of network invariants among alpha, beta, delta, and Omicron has been studied also in [38]. The presented results can also be considered as further extension of previous results presented in [38] in two ways. First, we confirm that the difference is not limited to network invariants, but also other protein structural measures, i.e. polarity and TM-Score confirm that Omicron is radically different from the other ones.

We also explore patterns of changes in a temporal dimension and compare the cumulative distribution of vaccination with the characteristics of the variant.Although we cannot infer any causality regarding the presence of vaccination as a force driving the evolution, we should note that the presence of vaccinations in a timeline is located in the middle of the first variants of SARS-CoV-2 and Omicron. Considering also the clinical characteristics of Omicron in terms of vaccine escape and neutralization of immune response, we can assume that the effect of all changes of Omicron may be related to the previous effects.

Finally, as illustrated in Figs. 13 and 14 we report that significant changes that appeared in the Omicron variants are subsequent to the increase in the number of vaccinations. We are aware however, that this does not imply any causal relation [39, 40]. Despite this we believe that more work in future should be done to shed out light on the relation between the vaccination campaign and the evolution of the SARS-CoV-2 virus.

**Fig 13.**
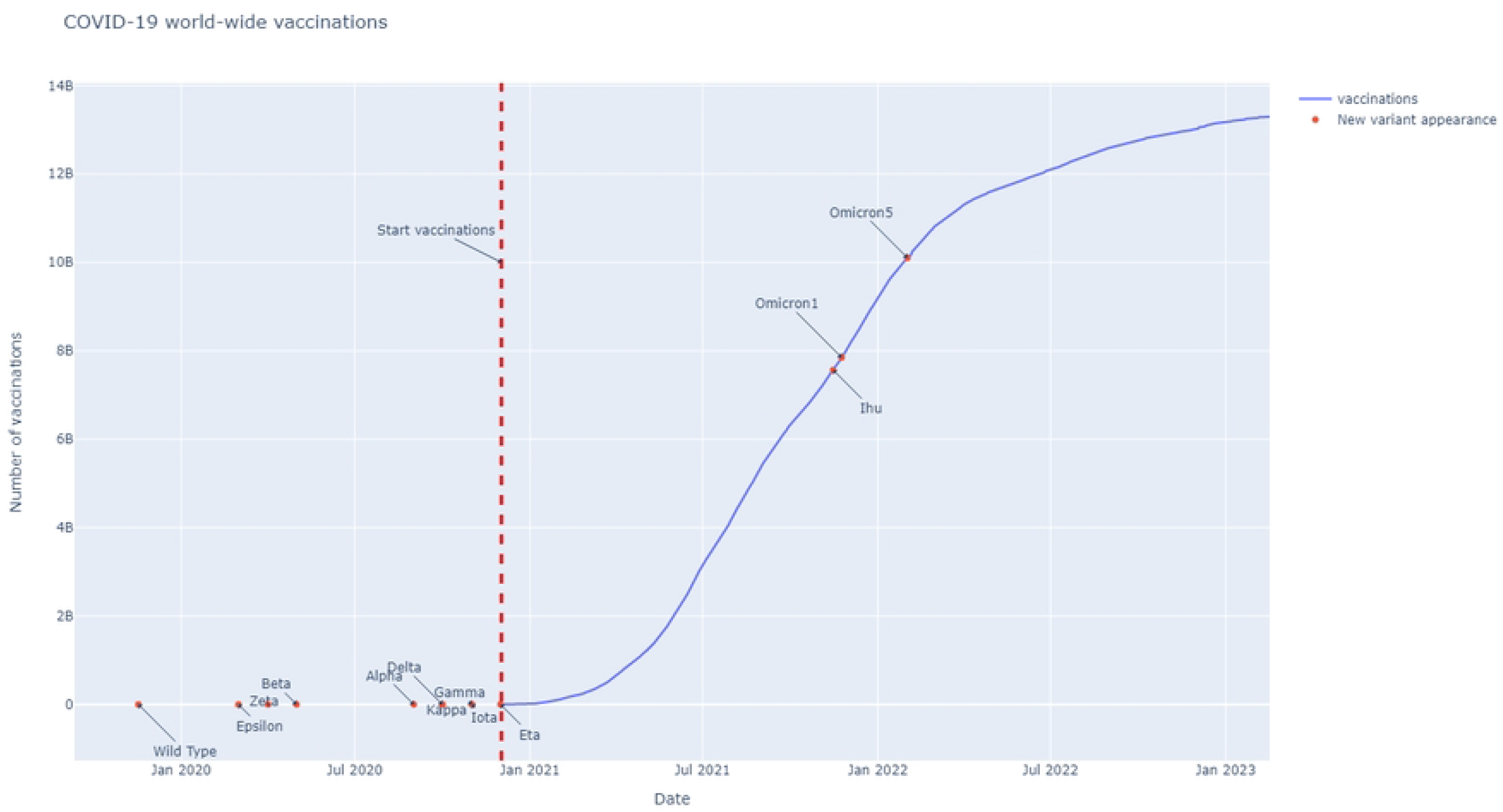
Covid-19 worldwide number of vaccinations and variants studied in this work. Variants are reported as red points on their first detection date.

**Fig 14.**
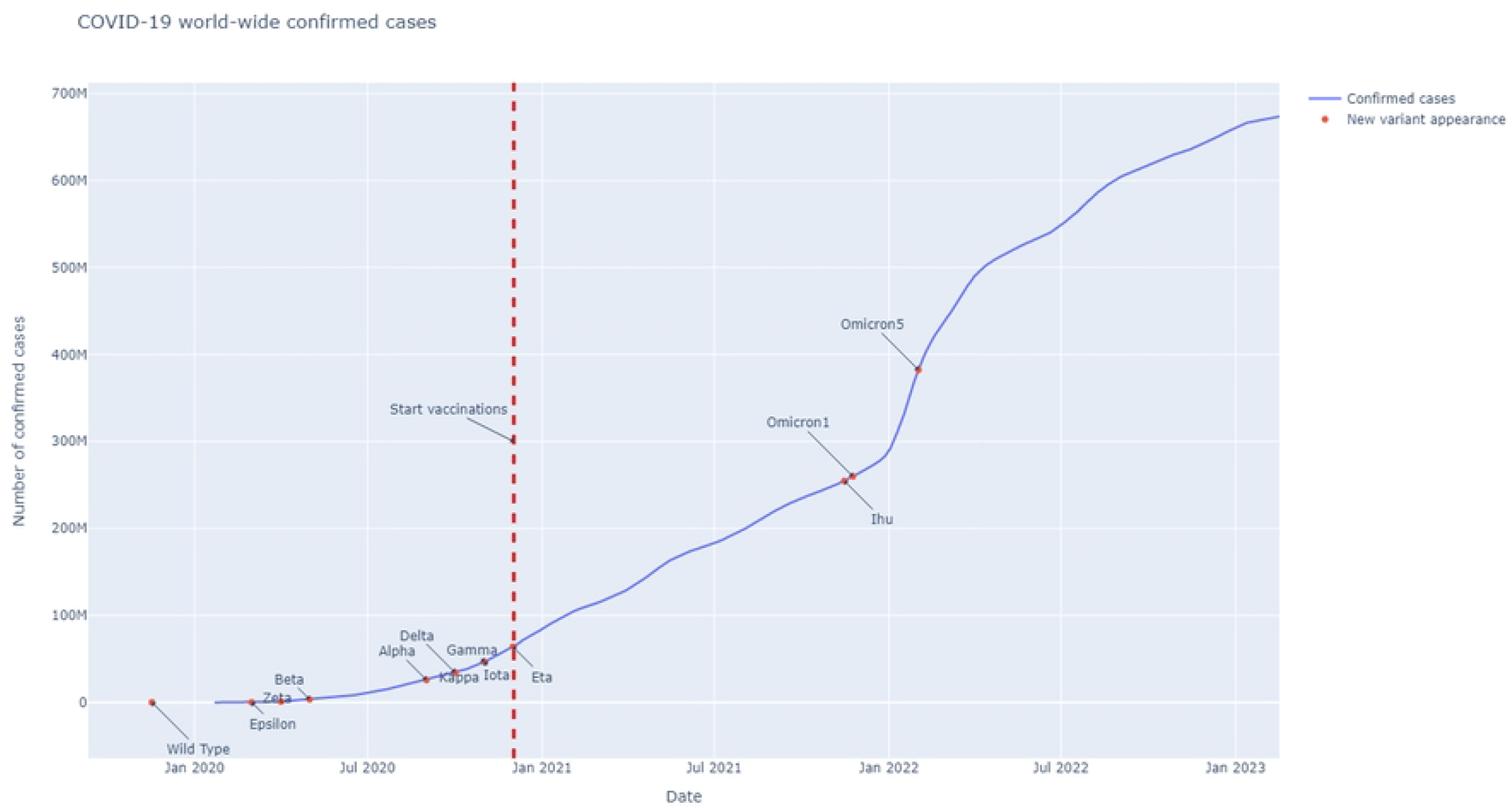
Covid-19 worldwide number of confirmed cases and variants studied in this work. Variants are reported as red points on their first detection date.

## Conclusion

In this work we have presented a pipeline of analysing mutations of the SARS-CoV-2 genome, focusing on the impact of the mutations on Spike protein. We analysed such changes from a timeline perspective. We observed that the Omicron variant presents significant changes with respect to the previous ones and Omicron appeared in parallel with the world wide vaccination campaign. Omicron’s structure presents many changes in the RBD domain and many mutation fall within the same structural region. Therefore, as a final remark, we would argue that further studies should focus on the possible relationship between the vaccination campaign and the appeareance of Omicron.

## Supporting information

The online git repository available at https://github.com/UgoLomoio/SARSCoV2_variants_PCN contains code, plots and datacx.

## Supplementary Figures

**Supp. Fig. 1.**
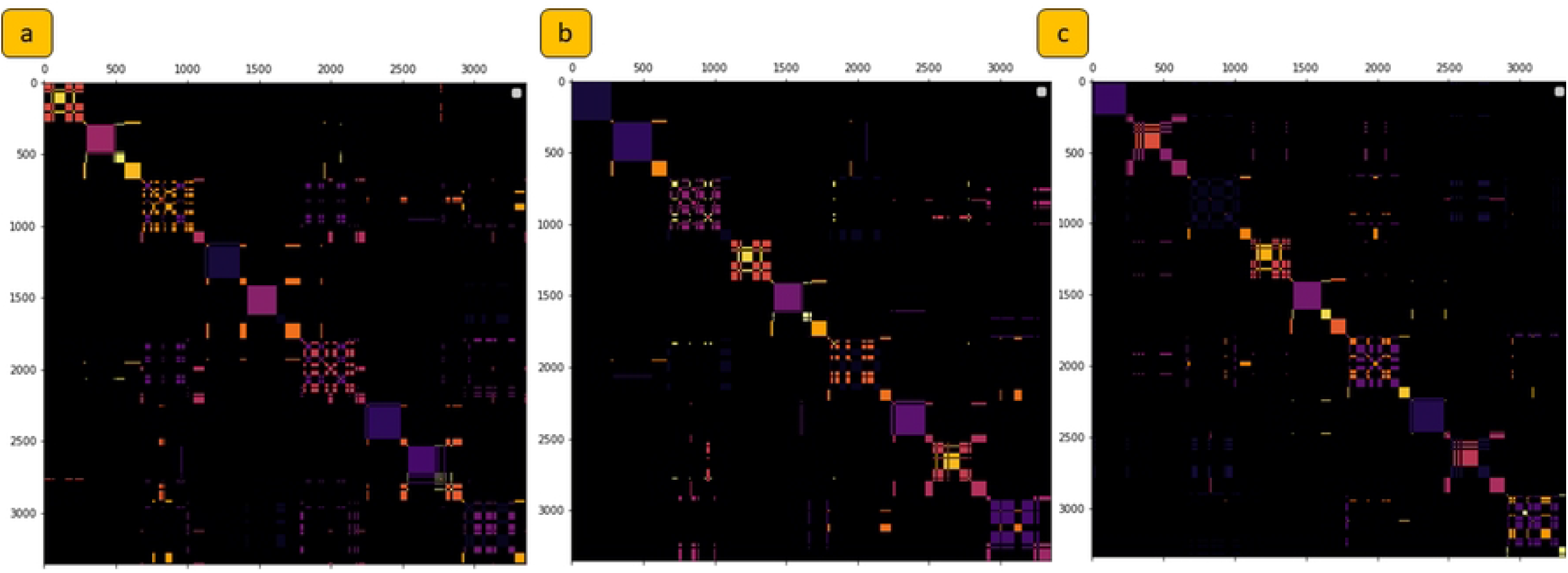
Clustermap plot for a) Wild Type, b) Delta, and c) Omicron_1_ variant. In a clustermap plot, communities are mapped in a matrix and visualized by means of heatmap.

## Acknowledgments

U.L., B.P., and P.H.G. are with the Department of Surgical and Medical Sciences, Magna Graecia University of Catanzaro.

G.T. is with e-Campus University, Novedrate Italy.

P.V. is with Dept. of Computer, Modelling, Electronics and System Engineering, University of Calabria.

PHG, PV, BP, and UL were partially funded by the MISE PON-VQA project. UL and BP Ph.D. fellow is partially supported by Relatech S.p.A.

